# Population Dynamics and Neuronal Polyploidy in the Developing Neocortex

**DOI:** 10.1101/2020.06.29.177469

**Authors:** Thomas Jungas, Mathieu Joseph, Mohamad-Ali Fawal, Alice Davy

## Abstract

The mammalian neocortex is composed of different subtypes of neurons which are generated during embryogenesis by sequential differentiation of neural progenitors. While molecular mechanisms that control neuronal production in the developing neocortex have been extensively studied, the dynamics and absolute numbers of the different progenitor and neuronal populations are still poorly characterized. Here we describe a medium throughput approach based on flow cytometry and well known identity markers of cortical subpopulations to collect quantitative data over the course of mouse neocortex development. We collected a complete dataset in a physiological developmental context on two progenitor and two neuron populations, including relative proportions and absolute numbers. Our study reveals unexpected numbers of progenitors. In addition, we discovered that a fraction of neurons in the developing mouse neocortex are polyploid.

## INTRODUCTION

In mammals, the brain stands apart from other organs due to its high degree of cellular diversity (Lodato et al., 2015) but also because the production of its functional units, neurons, is accomplished during pre-natal stages and mostly completed at birth. Indeed, postnatal expansion of brain size is mainly driven by an increase in non-neuronal cell populations and by neuronal maturation such as arborization, while generation of new neurons is limited after birth, especially in species with larger brain (Paredes et al., 2016). Consequently, neuronal production must be finely tuned during prenatal development in order to achieve species-specific stereotypical proportions of neuronal subtypes. It is now well accepted that changes in the process of neuronal production underlies brain evolution (Lui et al., 2011) and diseases (Juric-Sekhar and Hevner, 2019).

Mechanisms that control neuronal production in the developing brain have been extensively studied in the mouse neocortex and more recently in the human neocortex, leading to a general framework for this process in all mammals (Adnani et al., 2018; Miller et al., 2019). Briefly, radial glial cells, which are neural progenitors in the neocortex (also called apical progenitors) (Malatesta et al., 2000; Miyata et al., 2001; Noctor et al., 2001), give rise to all subtypes of projection neurons by direct or indirect neurogenesis, the latter process involving the production of intermediate progenitors also called basal progenitors (Borrell, 2019; Haubensak et al., 2004). One special feature of neuronal production in the mammalian neocortex is that the different subtypes of projection neurons are produced in a sequential manner, with deep layer neurons produced first and upper layer neurons produced last (Adnani et al., 2018).

Past studies have defined the temporal windows of production of each neuronal subtype as well as molecular mechanisms that control specific aspects of neurogenesis in the neocortex (Adnani et al., 2018; Greig et al., 2013). Yet, a gap of knowledge remains concerning neuronal and progenitor numbers as well as population dynamics in the developing neocortex. For instance, absolute numbers of neurons in the developing neocortex are usually referred to in vague terms such as “millions of neurons are produced” while absolute numbers of progenitors are usually ignored. Further, how these numbers evolve during neocortex development is not well characterized, despite the fact that this an important parameter to fully understand this developmental process. Given the growing interest in mathematical and computational studies that aim at providing predictive models for neocortex development and evolution (Cahalane et al., 2014; Caviness et al., 1995; Freret-Hodara et al., 2017; Llorca et al., 2019; Picco et al., 2018; Postel et al., 2019), addressing this quantitative gap is a timely and necessary challenge.

In the majority of published studies, neuron and progenitor numbers in the developing neocortex are collected using a time consuming process involving tissue processing and sectioning, followed by immunostaining with identity markers, imaging and manual counting within 2D regions of interest (ROI). Because these ROI represent only a tiny fraction of the tissue and neurogenesis is not synchronized throughout the tissue but progresses in a regionalized pattern (Dominguez et al., 2013), it is extremely difficult to extrapolate 3D cell numbers at organ level from these data. In contrast, flow cytometry allows automated analyses of entire organs, at single cell resolution and using several criteria. While the first flow cytometer was designed more than 50 years ago (Fulwyler, 1965), the use of flow cytometry to estimate cell numbers in the brain has been reported in only one study, at the adult stage (Young et al., 2012). Here we used a medium throughput approach based on flow cytometry and cortical sub-population identity markers commonly used in histology analyses to collect quantitative data on 2 progenitor and 2 neuron populations over the course of mouse neocortex development. This allowed us to establish the dynamics of cortical populations based on large number of counted cells and samples and to calculate the absolute number of the 4 cell types at 7 developmental stages. In addition, our simultaneous analysis of DNA content provided quantitative data on cell cycle parameters and led us to discover the presence of polyploid neurons in the developing mouse neocortex, notably in layer V.

## RESULTS

### A medium throughput pipeline to analyze neocortex samples by flow cytometry

To obtain a quantitative description of population dynamics in the developing neocortex, we focused on 4 main populations: 2 progenitor populations (apical and basal progenitors) and 2 neuronal populations (early born and late born neurons). Pax6 was used as a marker of apical progenitors (AP) and Tbr2 was used as a marker of basal progenitors (BP), Ctip2 was used as a marker of early born neurons and Satb2 was used as a marker of late born neurons (Figure 1A). We chose these transcription factors because they are amongst the most widely used identity markers in the field. Neocortical cells were collected from embryos at 7 developmental stages spanning corticogenesis, from E12.5 to E18.5. Briefly, the dorsal-most region of the telencephalon was dissected from each embryo and the tissue was dissociated mechanically. Following a filtration step, cells were fixed in 70% Ethanol and stored at −20°C (Figure 1B). Cells were immunostained with the different markers and with propidium iodide (PI) and samples were analyzed by flow cytometry using a 2-step gating strategy: a first morphometric-based gating on intact cells (P1) (Figure 1C) and a subsequent stringent gating based on PI incorporation (P2) to confirm debris and clumps exclusion and to eliminate cell doublets and cells in cytokinesis (P2 represents approximately 85% of cells gated in P1; see also Methods and (Gasnereau et al., 2007). Lastly, the proportion of each cell type in the P2 gated population was determined (Figure 1C).

**Figure 1:**
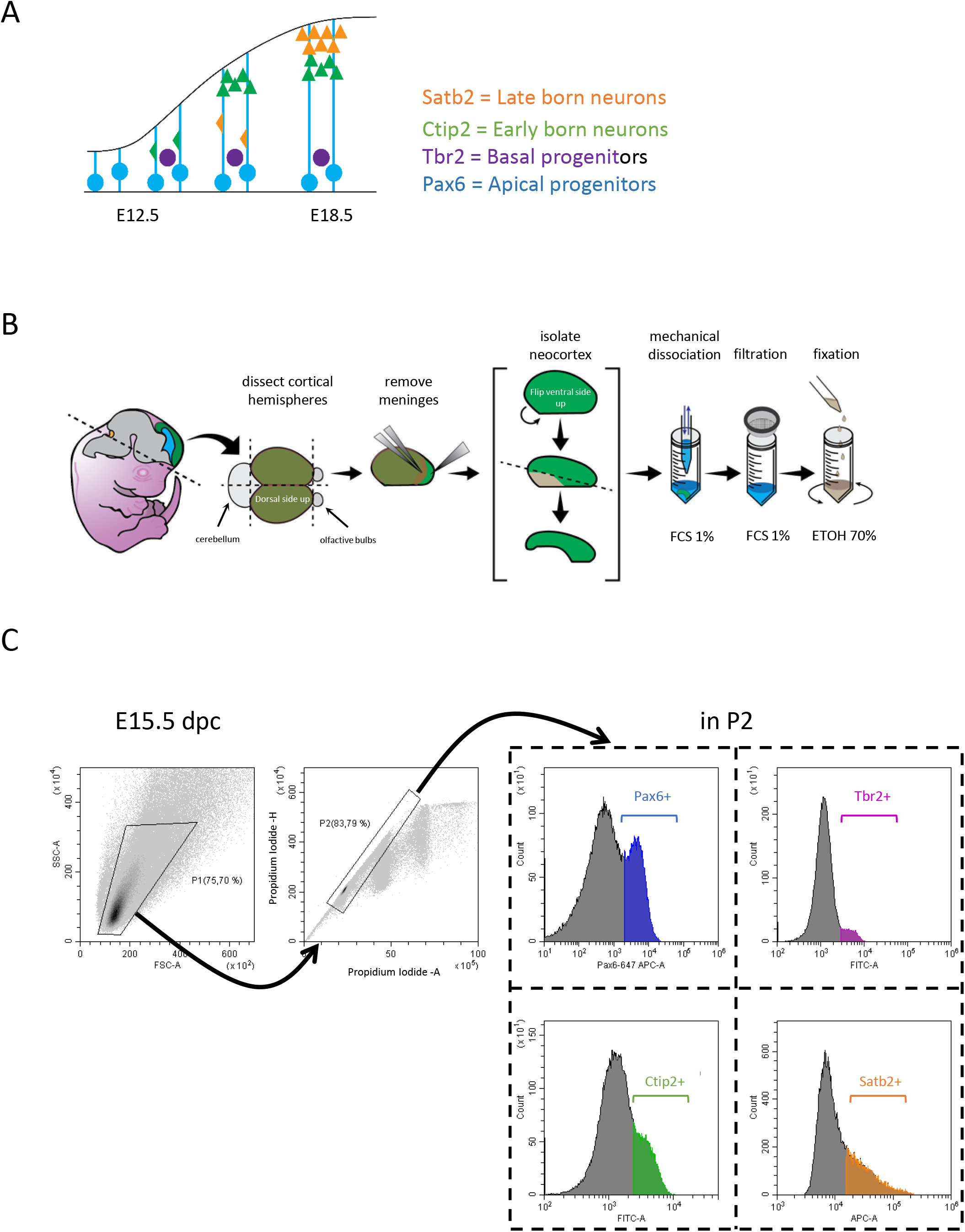
Description of the analytic pipeline. A: Schematic representation of mouse embryonic neocortex development with representation of 4 cell populations. Apical progenitors are in blue (Pax6+), Basal progenitors are in purple (Tbr2+), Early born neurons in green (Ctip2+), Late born neurons in orange (Satb2+). B: Experimental pipeline used to collect and prepare cortical cells from mouse embryos. C: Flow cytometry gating strategy. Representative Side scatter (SSC) versus Forward Scatter (FSC) density plot of all acquired events corresponding to a E15.5 embryo. The P1gate is shown (Upper left panel). Representative density plot for Propidium Iodide Height signal intensity (PI-H) versus Propidium Iodide Area signal intensity (PI-A) of P1 gated events showing the gate P2 corresponding to the singlets retained for further analysis (Upper second panel). Representative histogram plots corresponding to P2 gated events of one E15.5 embryo: Pax6 associated Alexafluor 647 signal intensity detected on APC-A channel (Pax6+ cells are in blue); Tbr2 associated Alexafluor 488 signal intensity detected on FITC-A channel (Tbr2+ cells are in purple); Ctip2 associated Alexafluor 488 signal intensity detected on FITC-A channel (Ctip2+ cells are in green); Satb2 associated Alexafluor 647 signal intensity detected on APC-A channel (Satb2+ cells are in orange).

Between 6-14 embryos from at least 2 different litters were analyzed by flow cytometry at each developmental stages for each marker. These analyses show that at early to mid-stages (E12.5-E14.5) of corticogenesis, the vast majority of neocortical cells are Pax6+ (Figure 2A). Between E14.5 and E15.5 we observe a sharp drop in the fraction of Pax6+ cells which is further diminished at E16.5 and reaches its lowest value at E18.5. The proportion of Tbr2+ cells remains low compared to Pax6+ cells, in the range of 10% of total cells from E12.5 to E14.5 (Figure 2B). From E15.5 to E18.5, this proportion decreases gradually (Figure 2B). Ctip2+ early born neurons are detected as early as E12.5 and their proportion increases sharply from E12.5 to E14.5, reaching its peak at E15.5 before dropping to a constant proportion between E16.5 and E18.5 (Figure 2C). Lastly, the proportion of Satb2+ cells shows a rapid growth phase between E14.5 and E18.5 reaching a maximum at E18.5 (Figure 2D).

**Figure 2:**
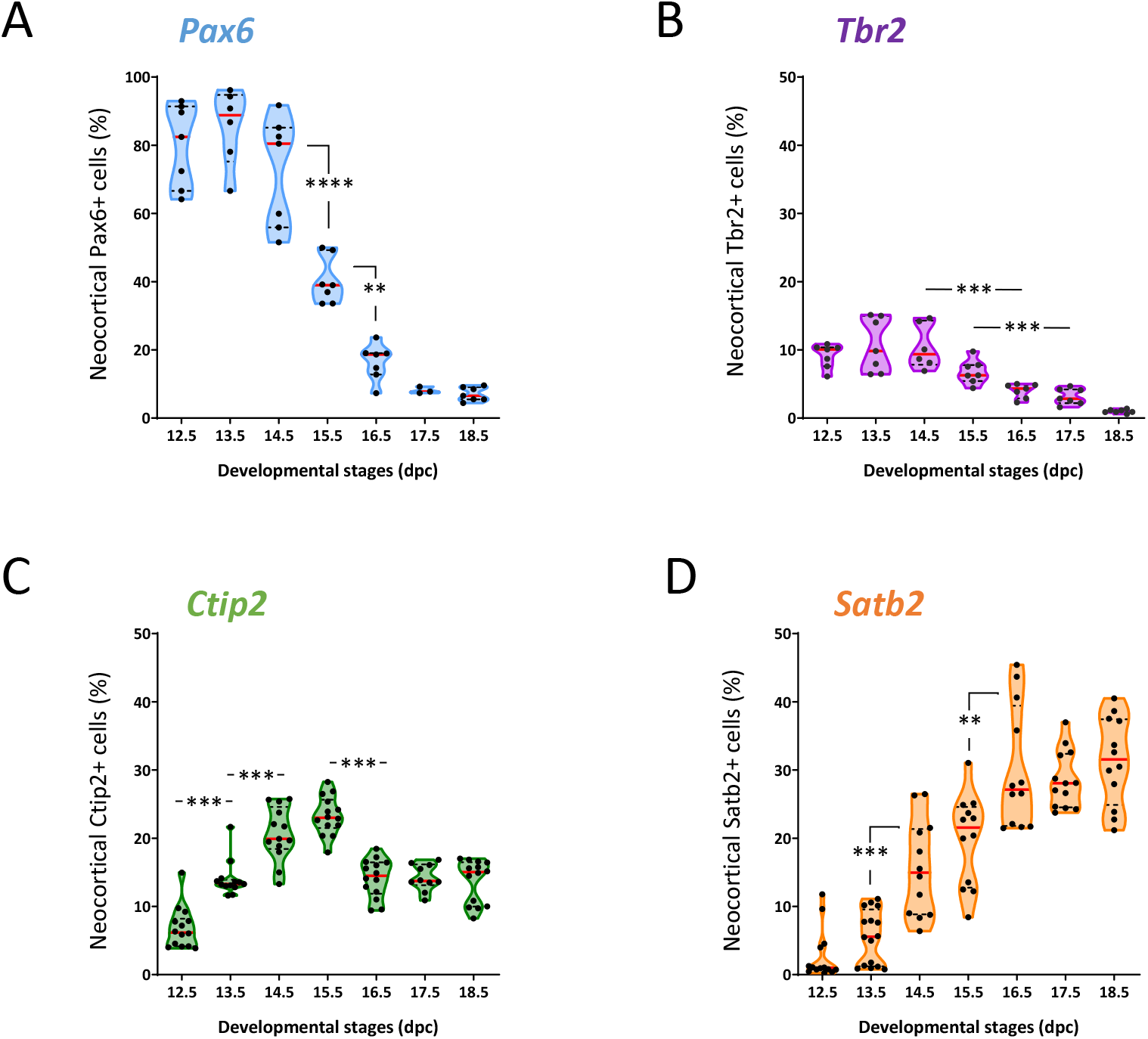
Evolution of the proportion of the 4 cell populations over the course of neocortex development. A-D: Relative proportion of indicated cell types at each indicated developmental stage. Each dot represents one embryo. Violin representation of the data displays the distribution (shape); the median (red line) and 1^st^ and 3^rd^ quartile (dotted black lines). One-way ANOVA with Tuckey’s post-hoc analysis was used for statistical analyses.

A more detailed analysis of the Pax6/Ctip2 data (both markers were analyzed together) reveals that at early stages of development a fraction of cells co-express both markers (Sup Figure 1A). Because these cells have a low Pax6 expression (the Pax6 intensity signal is shifted to the left), they might represent cells directly transitioning from AP to early born neurons, supporting the notion that some early born neurons are generated by direct neurogenesis (Sessa et al., 2008).

A similar analysis of the Tbr2/Satb2 data (both markers were analyzed together) did not reveal co-expression of both markers indicating sharp transition from progenitor to neuronal differentiation programs in BP (Sup Figure 1B). We also quantified the proportion of Tbr1+ neurons using our pipeline since Tbr1 is another identity transcription factor that is frequently used in the field to label deep layer neurons (Sup Figure 2). We observed that the proportion of Tbr1+ neurons increases from E12.5 to E13.5, reaching a plateau between E13.5 and E15.5 and decreasing at later stages. Overall, the proportion of these neurons remains close to or below the 10% mark (Sup Figure 2B).

Altogether these data provide a dynamic view of progenitor and neuron relative populations in the developing neocortex.

### Absolute numbers of progenitor and neuron populations in the developing neocortex

Although the evolution of cell populations relative to each other as shown above is very informative, it could be misleading. For instance, for Pax6+ cells (Figure 2A), the data gives the impression that cells are massively eliminated from E16.5 onwards while they could just be diluted. In order to provide a complete depiction of population dynamics in the developing neocortex, we designed protocols to calculate the absolute cell number of each progenitor and neuron populations, defined by the 4 identity markers. Two distinct protocols were used (Figure 3A). The first protocol makes use of beads that are mixed with the samples at a defined concentration (Ou et al., 2017). During flow cytometry analysis, the beads are detected by the flow cytometer and quantified. This provides a reference to extrapolate the volume of sample analyzed and subsequently, to calculate the absolute number of cells in each sample (see Methods section). However, newer generation flow cytometers directly measure the volume of sample analyzed which allows a direct estimation of absolute cell numbers as described in the second protocol (Figure 3A). We analyzed samples at the 7 developmental stages using both methods (Figure 3B, C). Comparison of cell numbers obtained with both methods for the same samples shows a high degree of correlation (Figure 3D) indicating that they can be used indiscriminately. Using both methods, we calculated that the total number of cells present in the developing neocortex varies from 1.23 10^5^ cells at E12.5 to 3.75 10^6^ cells at E18.5 which represents a 30-fold increase (Figure 3E). We also calculated total cell numbers in the adult neocortex which is between 21-24 10^6^ cells (Sup Figure 3A) representing a 5.8-fold increase. Next, we calculated the absolute number of each progenitor and neuron population throughout neocortex development. The number of Pax6+ cells sharply increases from E12.5 to E14.5 and steadily decreases between E15.5 and E17.5 (Figure 3F). This profile is consistent with the fact that at early stages, Pax6+ progenitors undergo both proliferative and neurogenic divisions, while at later stages neurogenic divisions are dominant thus depleting the pool. This profile also reveals that the absolute number of Pax6+ cells remains substantial at late stages (approximately 2.5 10^5^ cells at E18.5) contrary to what the relative data in Figure 2A could have suggested. The number of Tbr2+ progenitors builds up from E12.5 to E14.5, reaching a plateau of approximately 10^5^ cells and dropping at E18.5 (Figure 3G). This dynamic supports the fact that Tbr2+ basal progenitors contribute to neurogenesis mostly at later stages and are thus depleted at late stages. The number of Ctip2+ neurons steadily increases from E12.5 to E15.5 and remains fairly constant after E15.5 (Figure 3H), which is expected since Ctip2+ neurons are early born neurons. The number of Satb2+ neurons sharply rises from E14.5 to E18.5 with a steep slope (Figure 3I). The profile shows that the majority of Satb2+ neurons are generated after E14.5, consistent with the fact that Satb2+ neurons are late born neurons. Overall, absolute numbers for the 4 populations of progenitors and neurons show expected dynamics (Figure 3J). Yet our data reveals that the number of Tbr2+ progenitors at tissue level remains low compared to Pax6+ progenitors throughout corticogenesis (Figure 3J), which was not expected from published 2D data.

**Figure 3:**
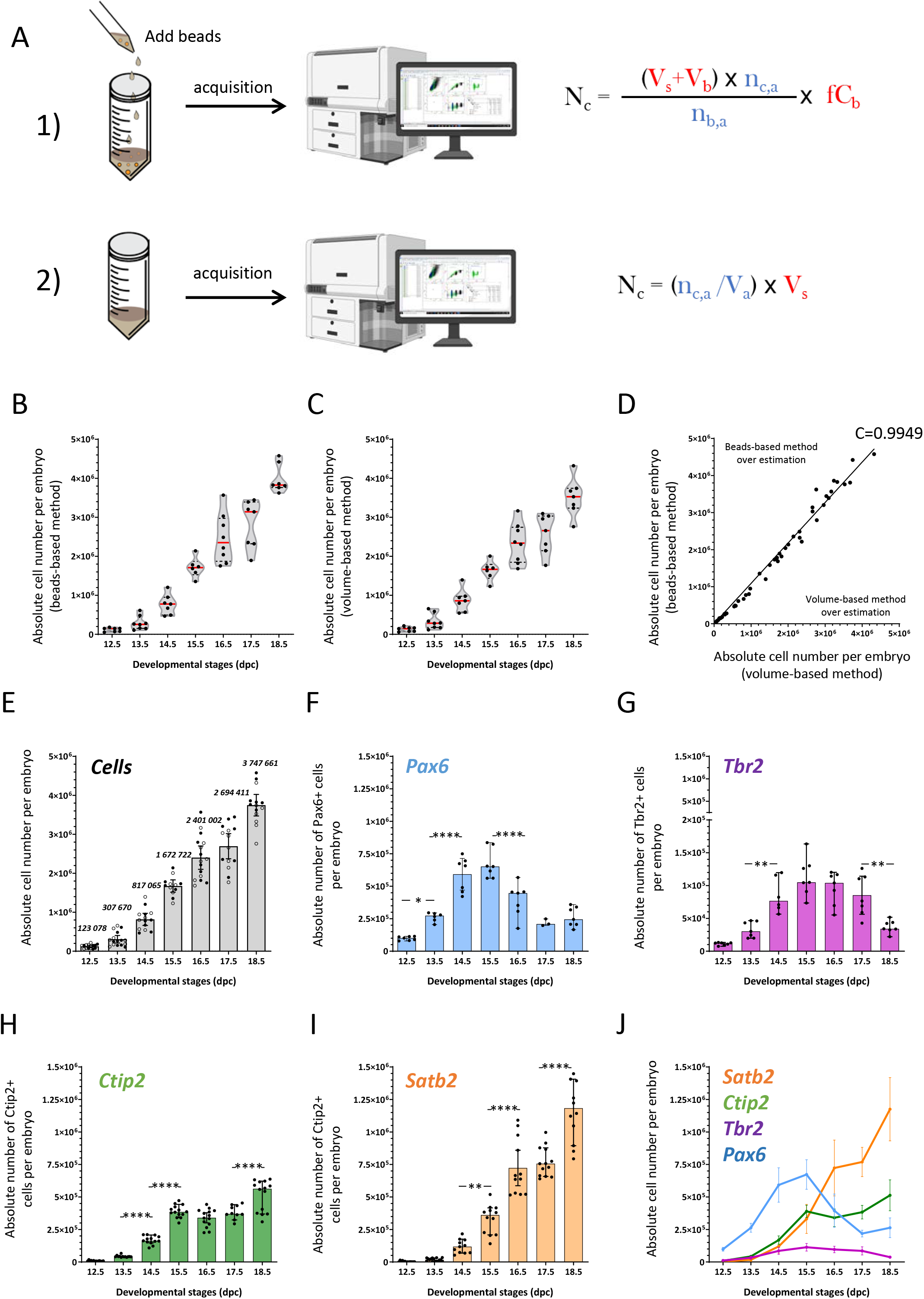
Evolution of absolute numbers of cells over the course of neocortex development. A: Schematic representation of the 2 procedures used to calculate absolute numbers. <u>Procedure 1</u>: calibrated fluorescent beads are mixed with the sample just before acquisition. Total cell number per embryo (N_c_) is calculated as indicated. Terms in red have to be collected prior to the experiment, terms in blue are given by the cytometer (upper panel). <u>Procedure 2</u>: This method needs specifically equipped flow cytometer that measure the volume of sample analyzed. Total cell number per embryo is calculated as indicated. Terms in red have to be collected prior to the experiment, terms in blue are given by the cytometer (lower panel). N_c_ is for total neocortical cell number per embryo, V_s_ is for initial volume of sample, V_b_ is for volume of beads added to sample, n_c,a_ is for number of cells acquired, fC_b_ is for final concentration of beads, nb,a is for number of beads acquired and V_a_ is for volume acquired. B: Total cell number in the neocortex at each developmental stage using the beads-based method. Each dot represents one embryo (also analyzed in C). Violin representation of the data displays the distribution (shape); the median (red line) and 1^st^ and 3^rd^ quartile (dotted black lines). C: Total cell number in the neocortex at each developmental stage using the volume-based method. Each dot represents one embryo (also analyzed in B). Violin representation of the data displays the distribution (shape); the median (red line) and 1^st^ and 3^rd^ quartile (dotted black lines). D: Comparison of the two procedures for the 55 embryos doubly analyzed. Each dot represents values for one embryo acquired with beads-based procedure (vertical axis) and volume-based procedure (horizontal axis). The correlation coefficient calculated with the spearman method is 0.9949 with a 95% confidence interval (95% CI) between 0.9908 and 0.9972 and a P value <0.0001. The line on the graph represent a simple linear regression with a slope of 1.091 (Y= 1.091X +0.000) with a 95% confidence interval between 1.067 and 1.115 and a P value <0.0001. E: Absolute number of cells calculated for each embryo with the beads-based method (filled dots) and the volume-base-method (empty dots). The histogram bars correspond to the mean (italic) with 95% confidence interval error bars of beads-based method values and volume-based method values collected at each developmental stage. F-I: Absolute number of indicated cell populations in the neocortex per embryo at each developmental stage. The histogram bars correspond to the mean with 95% confidence interval error bars. Each dot represents the calculated value for one embryo. J: Comparative dynamics of the absolute number of the 4 cell populations over the course of neocortex development: Pax6+ (blue), Tbr2+ (purple), Ctip2+ (green), Satb2+ (orange). Dots represent mean values for each developmental stage and error bars are for standard deviations to the mean. One-way ANOVA with Tuckey’s post-hoc analysis was used for statistical analyses.

Indeed, to compare our data to previously published data collected by counting cells on tissue section (2D method), we surveyed 7 previously published studies (Alsiö et al., 2013; Fei et al., 2014; Kischel et al., 2020; Knock et al., 2015; Lanctot et al., 2017; Postel et al., 2019; Wang et al., 2016; Yoon et al., 2017) and collected reported cell numbers at the 7 developmental stages (Sup Table 1). Graphical representation of the data (Sup Figure 3B) shows overall conservation of the profiles for each of the 4 cell populations with some differences, specifically in the number of Tbr2+ progenitors.

Altogether these results indicate that flow cytometry is a valid and rapid approach to calculate absolute cell numbers in tissue samples and that it is complementary to the 2D method.

### Analysis of cell cycle parameters by flow cytometry reveals that a fraction of Ctip2+ neurons are polyploid

To further characterize the dynamics of progenitor populations in the developing neocortex, we analyzed cell cycle parameters in Pax6+ and Tbr2+ progenitors, using PI staining. These analyses indicate that between 6 – 12 % of Pax6+ cells are in S-phase with a higher fraction at early stages and a lower but roughly stable fraction at later stages (Figure 4A, B). This may reflect the fact that cell cycle duration in AP lengthens at later stages (Calegari et al., 2005). The fraction of Tbr2+ cells in S-phase follows the same trend as Pax6+ cells, however, it is lower since it varies between 2.5-5 % (Figure 4C, D) which could explain the small number of Tbr2+ progenitors.

**Figure 4:**
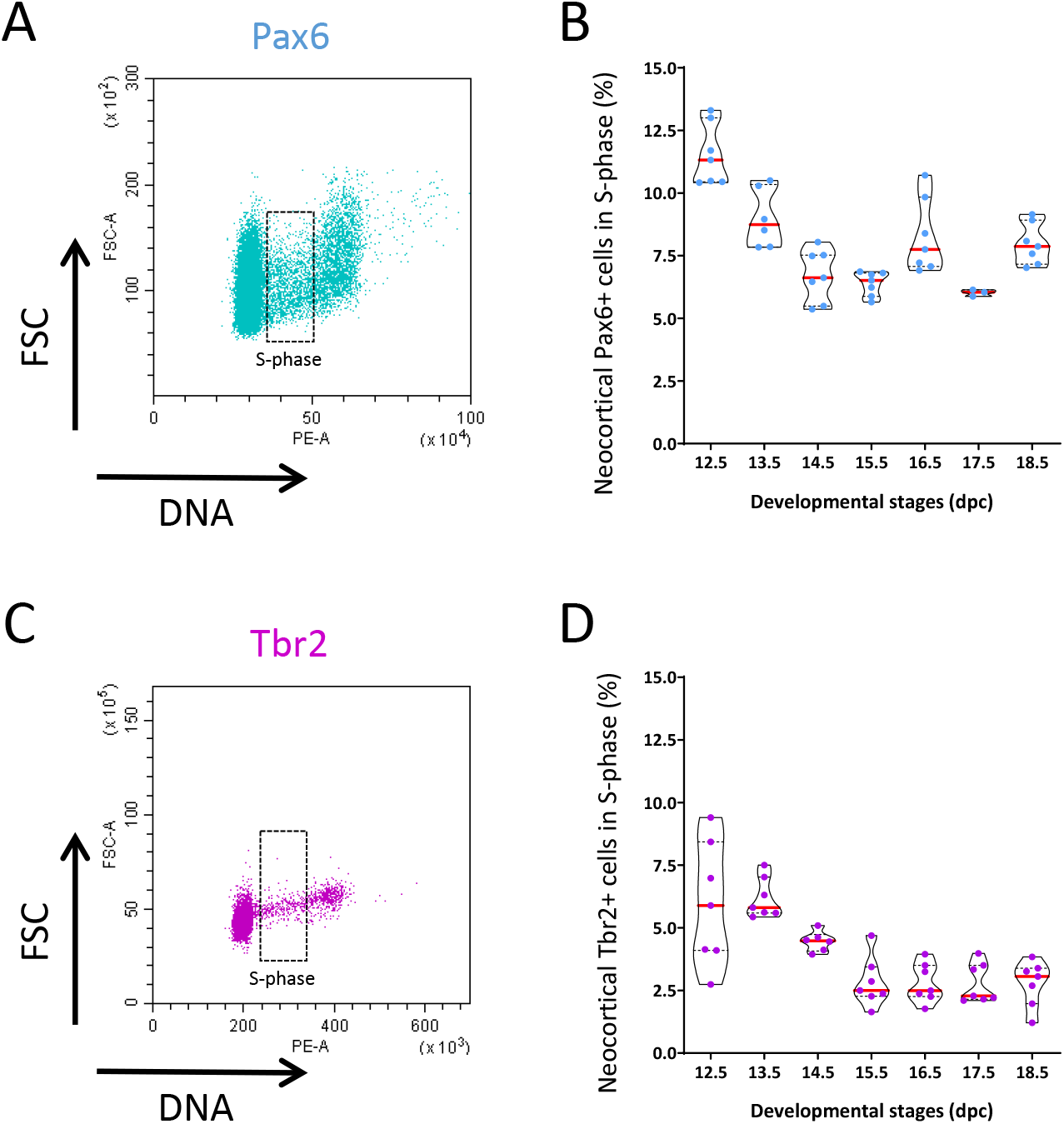
Cell cycle parameters for the 2 subtypes of cortical neural progenitors over the course of neocortex development. A: Representative dot plot of DNA content for Pax6+ cells estimated by Propidium Iodide incorporation (horizontal axis) versus FSC relative cell size (vertical axis) for one E15.5 embryo. Cells in S-phase are highlighted. B: Distribution of the proportion of Pax6+ cells in S-phase at each developmental stage. Each dot represents one embryo. Violin representation of the data displays the distribution (shape); the median (red line) and 1^st^ and 3^rd^ quartile (dotted black lines). C: Representative dot plot of DNA content for Tbr2+ cells estimated by Propidium Iodide incorporation (horizontal axis) versus FSC relative cell size (vertical axis) for one E15.5 embryo. Cells in S-phase are highlighted. B: Distribution of the proportion of Tbr2+ cells in S-phase at each developmental stage. Each dot represents one embryo. Violin representation of the data displays the distribution (shape); the median (red line) and 1^st^ and 3^rd^ quartile (dotted black lines). One-way ANOVA with Tuckey’s post-hoc analysis was used for statistical analyses.

We also analyzed DNA content of Ctip2+ and Satb2+ cells which are considered mostly post-mitotic. Unexpectedly, this analysis revealed that, despite very low numbers (below 2% throughout neocortex development), Ctip2+ and Satb2+ cells with a DNA S-phase like content could be detected at all stages (Sup Figure 4A-D) and more intriguing, Ctip2+, and to a lower extent Satb2+ cells, with a 4C DNA content could be detected (Figure 5A and Sup Figure 4C). A detailed analysis of the population of 4C Ctip2+ cells at all developmental stages reveals that it ranges between 2-5% and that it is maintained at late stages (Figure 5B). In terms of absolute numbers, 4C Ctip2+ cells represent close to 25.10^4^ cells in the neocortex at E18.5 (Figure 5C). To ensure that 4C Ctip2+ cells are postmitotic neurons and not progenitors transitioning to differentiation but still cycling, we performed a series of experiments using NeuN, a late marker of neuronal differentiation (Gusel’nikova and Korzhevskiy, 2015). We show that Ctip2+ neurons also express NeuN and that 4C NeuN+ neurons can be detected in the developing neocortex (Figure 5D). Further, NeuN+ 4C neurons are enriched for Ctip2+ at later stages of development (Figure 5E). Ctip2 is a marker of early born layer V neurons and it has been shown previously in the adult rat neocortex that some layer V neurons are polyploid (Sigl-Glockner and Brecht, 2017). To test whether 4C neurons could be polyploid, we first confirmed that our flow cytometry pipeline is efficient at detecting polyploid cells by processing adult liver tissues as positive controls. The flow cytometry pipeline was able to detect polyploid cells in the liver, with a higher accumulation in older tissues as expected (Chao et al., 2017) (Sup Figure 5A, B). Next, we directly assessed chromosomal content in 4C neurons using DNA FISH (fluorescent in situ hybridization) with locus-specific chromosomal probes. Dissociated cells from neocortical tissue at E18.5 were incubated with NeuroFluor^™^ NeuO membrane-permeable fluorescent probe which labels live neurons and Vybrant^®^ DyeCycle^™^ Ruby Stain for DNA content quantification. Cells were then FACS-sorted based on DNA content in 2C versus ≥4C cells and based on neuronal (NeuO+) versus non neuronal (NeuO-) cells (Figure 5F). Sorted cells were analyzed by FISH using probes for 2 sets of chromosomes. 2C cells typically had 2 red spots and 2 green spots, as expected (Figure 5F). On the other hand, ≥4C cells had more than 2 spots of each color (Figure 5F), indicating an increased number of chromosomes which is consistent with polyploidy in post-mitotic cells.

**Figure 5:**
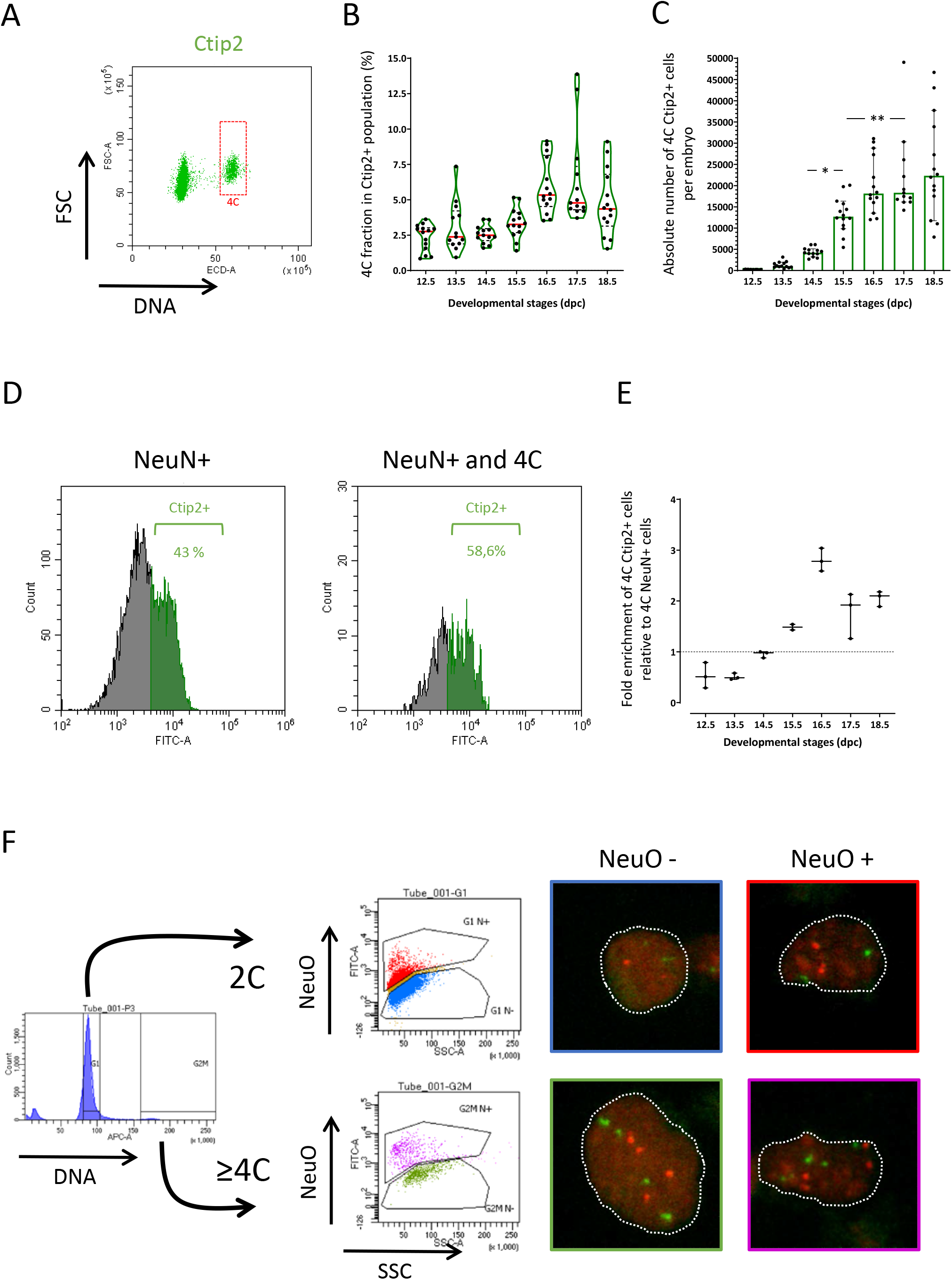
Identification of polyploid neurons in the developing neocortex. A: Representative dot plot of DNA content for Ctip2+ cells estimated by Propidium Iodide incorporation (horizontal axis) versus FSC relative cell size (vertical axis) for one E15.5 embryo. Cells with a 4C DNA content are highlighted. B: Distribution of the proportion of Ctip2+ cells with a 4C DNA content at each developmental stage. Each dot represents one embryo. Violin representation of the data displays the distribution (shape); the median (red line) and 1^st^ and 3^rd^ quartile (dotted black lines). C: Absolute number of Ctip2+ cells with a 4C DNA content in the neocortex per embryo at each developmental stage. The histogram bars correspond to the mean with 95% confidence interval error bars. Each dot represents the calculated value for one embryo. D: Representative histogram plots for one E15.5 embryo highlighting Ctip2+ cells (green) within the NeuN+ fraction (left panel) and within the NeuN+ fraction with a 4C DNA content (right panel). E: Ratio of NeuN+/4C/Ctip2+ fraction versus NeuN+/Ctip2+ fraction at each developmental stage. Each dot represents one embryo and error bars are for 95% confidence interval. F: Dissociated neocortical cells from E18.5 embryo were FACS-sorted based on DNA content and on labelling with NeuroFluor^™^ NeuO. 4 distinct populations were collected based on the following parameters: 2C DNA content versus ≥4C DNA content; NeuO negative cells (non neurons) versus NeuO positive cells (neurons). Sorted cells were then processed for fluorescent in situ hybridization (FISH) using two DNA probes specific for the mouse chromosome 11 (red spots) and chromosome 2 (green spots). Dotted white lines highlight nuclei shape in representative images for each sorted populations. One-way ANOVA with Tuckey’s post-hoc analysis was used for statistical analyses.

Altogether these results reveal that polyploid neurons are present in the developing neocortex and that a significant fraction of these are Ctip2+ at late developmental stages. More generally, these results indicate that the flow cytometry pipeline is useful to identify uncharacterized cell populations in the developing neocortex.

## DISCUSSION

To fully understand neocortex development which is a highly dynamic process, it is necessary to collect and integrate quantitative data at all scales, from molecules to cell populations. The advent of single cell RNA sequencing (scRNA-Seq) technology in the last few years has led to an unprecedented drive to characterize cell populations based on their transcriptional profiles. This has been extremely informative in the field of developmental biology and more specifically for studies of lineage progression during organogenesis. For instance in the developing neocortex it was shown that dynamic waves of transcriptional programs drive neuronal differentiation and temporal patterning (Telley et al., 2019; Telley et al., 2016). Given that scRNA-Seq detects mRNA, flow cytometry represents a useful complementary approach that allows multiparametric analyses of large numbers of cells at single cell resolution, allowing the detection of proteins or DNA content (McKinnon, 2018). In addition, flow cytometry provides quantitative data at population level that is complementary to clonal analyses performed in the developing neocortex that provide quantitative information on clone size and composition (Beattie et al., 2020; Gao et al., 2014).

Here we report quantitative parameters on population dynamics in the developing neocortex, focusing on 2 progenitor populations and 2 neuron populations. We chose a combination of identity markers that label a majority of progenitors and projection neurons in the developing neocortex. (Woodworth et al., 2012). The first parameter we analyzed, which is commonly analyzed with flow cytometry, is the relative proportion of the 4 cell populations in the developing neocortex. Of note, similar estimates of relative proportions of progenitor versus neuron populations were reported very recently using different markers, tissue dissociation and manual counting (Sahara et al., 2020).

One of the real advantage of the flow cytometry pipeline described here is the possibility to extract absolute cell numbers for each population. Here we compared two methods, a beads-based method and a volume-based method which provided equivalent estimate of cell numbers. The first method, which is based on the use of calibrated beads at known concentration, is more accessible but requires rigor in the standardization and handling of the beads and can be affected by the presence of bead aggregates that distort the calculations (Ou et al., 2017). The second method requires a cytometer equipped with either a peristaltic pump or an injector sample. With this method, attention must be paid to the performance of the fluidic tubing (Brunck et al., 2014). With both methods we report total cell numbers in adult brain that correspond to numbers previously reported using a different method, the “isotropic fractionator” (Herculano-Houzel et al., 2013).

No data is currently available on absolute cell numbers in the developing neocortex so it is not possible to compare our cytometry pipeline to other approaches. However, we show that the dynamics of cell numbers for each population is overall similar to what has been reported using 2D manual counting, yet we also noted differences. For instance, at late stages of development Satb2+ cells show a drop in cell numbers in the 2D method, probably due to the fact that at late stages of development, cell density decreases due to neuronal arborization and thus the number of cells in a ROI decreases. A second difference is in the number of Tbr2+ cells which is lower in the cytometry data, perhaps reflecting the fact that 2D analyses usually focus on lateral regions of the neocortex while cytometry analyses the entire tissue. Our results suggest that these cells do not represent a population of typical transient amplifying intermediate progenitors, since their number remains low and only 2-5% of Tbr2+ cells are in S-Phase throughout neocortex development indicating that they are slowly dividing cells. Interestingly, asymmetric cell death of Tbr2+ daughter cells as well as division with delayed terminal differentiation was reported recently using clonal analysis in the mouse neocortex (Mihalas and Hevner, 2018), which might contribute to the observed dynamics of the Tbr2+ population.

We observed some heterogeneity in the data at each developmental stage with all the markers we used. This heterogeneity partly reflects the fact that we staged the embryos based on the day postcoitum and not based on developmental landmarks, which means that some embryos could have been slightly advanced or delayed with respect to the indicated stage. Yet, we cannot completely exclude that part of the heterogeneity is due to technical bias in the procedure for instance during the steps of dissection and dissociation. Nevertheless, collectively, this quantitative dataset provides a detailed description of the dynamics of the 4 populations both in terms of relative and absolute numbers and should be a useful reference for future analyses of genetic mutants or to calibrate mathematical and computational models of neocortex development as was described very recently (Sahara et al., 2020).

We identified an uncharacterized population of polyploid neurons in the neocortex at pre-natal stages. Unlike in the developing neocortex, the presence of polyploid neurons in the perinatal or postnatal rodent neocortex has been reported previously (Jungas et al., 2016; Lopez-Sanchez and Frade, 2013; Sigl-Glockner and Brecht, 2017) and strong evidence suggest that these neurons are located in layer V and are the largest neurons in the neocortex (Sigl-Glockner and Brecht, 2017). In humans, neuronal polyploidy in the cortex has been associated with pathological states such as Alzeihmer’s disease (Arendt et al., 2010; Yang et al., 2001), but this issue remains controversial (Westra et al., 2009). However, in physiological conditions, giant neurons discovered by Vladimir Betz are present in layer V of the human primary motor cortex (Kushchayev et al., 2012). Whether these neurons are polyploid is not known. Our data suggest that in physiological conditions, acquisition of the polyploid state might not be a late postnatal event, but rather an early event, at the time of, or shortly after, neuronal production. Whether polyploidy confers specific functions to these neurons is an exciting open question for the future.

## ACKNOWLEDGEMENTS

We are grateful to Christophe Audouard for his help with collecting samples and to Sophie Bel-Vialar and Alain Vincent for helpful comments on the manuscript. We acknowledge core support from the TRI-IPBS imaging platform for FACS sorting and from ANEXPLO/CBI for housing and caring for the mice. This work was funded by ANR (Agence Nationale de la Recherche) (ANR-15-CE13-0010-01).

## AUTHOR CONTRIBUTIONS

AD designed and supervised the project; TJ and MJ performed experiments; AD, TJ and MAF analyzed and interpreted the data; AD and TJ wrote the manuscript; MAF edited the manuscript.

## MATERIALS AND METHODS

### Animals

Mice were housed and bred at the CBI/Anexplo animal facility according to standard SPF procedures. Mice were on a mixed 129S4/C57Bl6JRj wild type genetic background. Timed-pregnant mice were sacrificed by cervical dislocation while embryos were euthanized by decapitation. These methods of euthanasia were approved by the appropriate Ethics Committee (APAFIS#1289-2015110609133558 v5). Both male and female embryos were collected and analyzed at 7 developmental stages from E12.5 to E18.5.

### Tissue collection

Tissue collection was performed in the early afternoon to minimize inter-litter variability. Time controlled pregnant female mice were euthanized by cervical dislocation and collected fetuses were immediately placed in ice cold PBS leading to hypothermic anesthesia before being euthanized by decapitation. The brain was first removed from the head cavity using forceps and olfactive bulbs were removed with forceps facilitating simultaneous removal of the meninges. To isolate the neocortex, the brain was placed ventrally on the dissection plate. Using oblique pressure with forceps, the ventral brain was eliminated. Then, hemispheres were positioned with their outer surface in contact with the bottom of the plate to visualize the ganglionic eminences which were removed with a scalpel. The 2 cortical hemispheres were then pooled and transferred into individual annotated tubes.

### Cell dissociation and immunostaining

Neocortical tissue transferred into individual tubes in ice cold 1% FCS-PBS solution were homogenized by mechanical dissociation using 1000μl tips until no aggregates were visible. To eliminate clumps incompatible with flow cytometry, the cell suspension was filtered through a 40 μm nylon mesh placed on a 50 ml conical tube using a 10 ml piston syringe and the filter was rinsed with at least 4 ml of 1% FCS-PBS solution. The cell suspension was transferred into a 15 ml conical tube and cells were centrifuged at 1500 rpm for 10 minutes at 4°C. To potentiate stoichiometric DNA staining and facilitate day by day embryos collection and long term storage, cells were immediately fixed using 70% ethanol. Briefly, the cell pellet was suspended in 1.2 ml of ice cold PBS using a 1000μl tips and 3 ml of 100% ethanol stored at −20°C was added dropwise on the cell suspension while vortexing. Samples were then stored at −20°C.

For immunostaining samples were centrifuged at 2200 rpm for 15 minutes and pellets were washed once with a large volume of ice cold PBS (10 ml) to eliminate ethanol traces. Primary antibodies were diluted in 200 μl/tube PBTA (1% FCS, 1% BSA and 0.1% Triton X-100 in PBS) with the adapted dilution factors (see STAR Method). Cells were incubated with primary antibodies overnight at 4°C. The next day, 10 ml ice cold PBS was distributed in each tube, pellets were suspended with manual shaking before a centrifugation step at 4°C. Cells were then incubated for 2 hours at room temperature in the dark with 200μl of diluted secondary antibodies. Cells were rinsed with 10 ml ice cold PBS and were resuspended with a volume of ready to use FxCycle^™^ PI/Rnase solution, according to the manufacturer’s instructions (Invitrogen, F10797).

### Primary and secondary antibodies

Antibodies were used at the following dilutions: rabbit anti-Pax6 (Biolegend, polyclonal, 1:300); Rat anti-CTIP2 (Abcam, clone 25B6, 1:300); Rat anti-tbr2 (Invitrogen, Eomes clone Dan11mag, 1:100); mouse anti-Satb2 (Abcam, clone SATBA4B10,1:50); Mouse anti-Neun (Millipore, clone A60, 1:200), Rabbit anti-Tbr1 (Abcam, polyclonal, 1:100). Anti-Rabbit, Mouse or Rat conjugated with Alexa Fluor 488 or Alexa Fluor 647 fluorophore secondary antibodies (Life Technologies) were used at 1:500 in PBTA solution.

### Flow cytometry analysis

Flow cytometry was performed using a Cytoflex-S Flow cytometer (Beckman Coulter Company) equipped with a 405nm Ultra violet laser, a 488 nm blue laser, a 561 nm yellow-green laser and a 640 nm red laser. Emission filters used were respectively BP 450/50 for Alexa Fluor 405 fluorochrome detection, BP 530/30 for Alexa Fluor 488 fluorochrome detection, BP 610/20 for Propidium iodide detection and BP 660/20 for Alexa Fluor 647 fluorochrome detection. Data was collected and analyzed using CytExpert software (Beckman Coulter Company) and displayed exponential scaling except for propidium iodide for whom linear scaling was used. For maximum doublets resolution, minimal flow rate (< 1000 events/second) was used for acquisition. A range of 20000 to 200000 events was acquired in each experiment.

For analysis, nuclei were gated as follow: A first clean-up gate was drawn on the forward (FSC-A) versus Side (SSC-A) scatter plot to eliminate clumps and debris. Cellular debris were also confirmed due to poor incorporation of DNA intercalating agent. Second, doublets were excluded by gating the DNA pulse area (PE-A) versus its corresponding height (PE-H). To set compensations when necessary and define positive signal during analysis - for each immunostaining – immunostained cells with single primary antibodies and all secondary antibodies were compared.

For cell cycle and ploidy analysis DNA content histograms were generated and G1/G0, S-phase and G2/M or tetraploid regions where manually determined according to Darzynkiewicz methodology (Darzynkiewicz et al., 2010).

To estimate absolute number using the beads-based methods, 40μl or 50 μl of 7 μm calibrated beads (thermo Scientific, Cyto-Cal^™^Count Control, FC07) were added immediately after PI resuspension. Beads were placed under agitation for at least 1 hour and vigorously vortexed prior to pipetting to eliminate clumps and doublets. The method described above is applicable to any flow cytometer, and we validated it on both Calibur (Becton Dickinson) and Cytoflex S (Beckman Coulter) machines.

### Absolute number calculations

For the beads-based method, the volume of beads added, their concentration and the total volume of cell suspension (corresponding to 2 neocortical hemispheres) have to be recorded for each embryo prior to analysis. At the end of flow cytometry analysis, the number of cells and the number of beads analyzed have to be recorded. The total number of cells in the sample is calculated as follows: (sample and bead volume x number of cells analyzed) / (final bead concentration / number of beads analyzed). For the volume-based method, only the total volume of cell suspension (corresponding to 2 neocortical hemispheres) has to be recorded for each embryo prior to analysis. The total number of cells in the sample is calculated as follows: (number of cells analyzed / volume analyzed) x volume sample. Total cell number at each developmental stage has been calculated for n=6-8 embryos with both methods combined and the mean of these 12-16 values has been used as a reference to estimate absolute numbers of each cell population as follows: mean absolute total number of cells x fraction (%) of cells positive for identity marker.

### Cell sorting

For cell sorting, cells from E18.5 dpc embryos were obtained as described above. Tissue from six embryos was pooled. Approximatively 10.10^6^ cells were incubated for 2 hours in DMEM/F12 (cat# 31331-08, Gibco) containing 30 % glucose (cat# UG3050, Euromedex) and 5 ng/ml FGF (cta# S029, Sigma-Aldrich) with 1:800 dilution of membrane permeant NeuroFluor^™^NeuO probe (cat# 01801, StemCell technologies) to detect neurons and 1:500 dilution of cell permeable DNA dye Vybrant^®^ DyeCycle^™^ Ruby Stain (cat# V10309, ThermoFischer scientific) to analyse DNA content. Unwashed samples were sorted on a FacsAriaIII cytometer equipped with the BD FACSDiva 8.0.2 version software. As described above, a first gate was applied to eliminate debris and clumps using FSC versus SSC parameters. A second and third gates were consecutively applied to eliminate remaining debris and cell doublets respectively based on FSC-height signal versus FSC-width signal and SSC-height signal versus SSC-width signal. NeuO positive threshold was determined using an aliquot of the cell suspension treated with the same conditions but without adding NeuroFluor^™^NeuO probe. 2C and 4C cell DNA content was determined based on Vybrant^®^ DyeCycle^™^ Ruby Stain fluorescence intensity with the principal first peak representing 2C fraction and the second detectable peak, at approximatively 2 fold the first peak value, representing 4C fraction. To limit sorting contamination of key populations, 4 ways sorting was performed as follow: NeuO+/2C at the far left outlet nozzle, NeuO-/4C at left outlet nozzle, NeuO+/4C at right outlet nozzle and NeuO-/2C at far right outlet nozzle. To preserve cell integrity and viability, cells were collected in tubes containing small volume of ice cold DMEM/F12 complete medium.

### DNA Fluorescent in situ Hybridization

Live sorted cells were transferred to 15 ml conical tubes and centrifuged for 15 minutes at 1500 rpm at 4°C. Pellets were suspended with 37°C PBS supplemented with 1% fetal calf serum at a concentration ranging from 10^4^ to 10^5^ cells/ml and distributed on Poly-L-Lysine (cat# P4707, Sigma-Aldrich) precoated coverslips for 3 hours at 37°C in 5% CO2 incubator to allow cells to adhere. Medium was replaced with 1ml of ice cold (3:1) methanol: acetic acid solution for 5 minutes at room temperature. Coverslips were rinsed twice with ice cold PBS and air dried. 10 μl drops of dual color directly-labeled DNA probes (Leica Biosystems, KBI-30501) to detect chromosomes 11 (11qE1) and 2 (2qH3) with respectively Platinum^™^Bright 550 fluorophore (red spots) and Platinum^™^Bright 495 fluorophore (green spots) were spotted on (3:1) methanol: acetic pre-cleaned slides. Coverslips were flipped over the drops with cells facing the liquid, sealed with removable silicone cement and immediately placed on a 75°C hotplate for 2 minutes to simultaneously denature probes and DNA. Samples were then incubated overnight at 37°C in the dark in a humid chamber. On the next day cement was gently removed and coverslips were dipped in 72°C pre-warmed 0.4X SSC for 2 minutes followed with a 30 seconds room temperature bath in 2X SSC/0.05% Tween 20. Cells on coverslips were counterstained with 1:1000 DAPI solution for 15 minutes at room temperature in the dark. Finally, coverslips were dipped in water, mounted on a glass slide with Mowiol before imaging on a confocal microscope (Leica SP8 inverted confocal microscope) equipped with Las X acquisition software (Leica microsystems). Images were analyzed with ImageJ (NIH).

### Quantification and statistical analyses

Quantitative data were collected from n=6-14 embryos for each developmental stage. Graphical representations of flow cytometry data were done using Prism8 GraphPad. All graphs show individual data points corresponding to individual embryos. Violins plot were used to emphasize sample distribution for each developmental stage. Each dot represents the measured value for one single embryo. Red line represents median of values and dotted black lines the 95 % confidence interval. Histogram bars were used to represent the mean of absolute number data with dots that represent the calculated value for one single embryo. Errors bars are for standard deviation to the calculated mean. On Figure 3D, dots correspond to calculated values for one single embryo with respectively the beads-based method (vertical axis) and the volume-based method (horizontal axis). The correlation index was calculated using the spearman method. Statistical analyses were performed with Prism8 GraphPad. One-way ANOVA with Tuckey’s post-hoc analysis was used for all statistical analyses. *p<0.05; **p<0.01; ***p<0.001; ****p<0.0001.

## REFERENCES

Adnani, L., Han, S., Li, S., Mattar†, P., and Schuurmans, C. (2018). Mechanisms of Cortical Differentiation. Int Rev Cell Mol Biol 336, 223–320.

Alsiö, J.M., Tarchini, B., Cayouette, M., and Livesey, F.J. (2013). Ikaros promotes early-born neuronal fates in the cerebral cortex. Proc Natl Acad Sci U S A. 110, E716–725.

Arendt, T., Bruckner, M.K., Mosch, B., and Losche, A. (2010). Selective cell death of hyperploid neurons in Alzheimer’s disease. Am J Pathol 177, 15–20.

Beattie, R., Streicher, C., Amberg, N., Cheung, G., Contreras, X., Hansen, A.H., and Hippenmeyer, S. (2020). Lineage Tracing and Clonal Analysis in Developing Cerebral Cortex Using Mosaic Analysis with Double Markers (MADM). J Vis Exp.

Borrell, V. (2019). Recent advances in understanding neocortical development. F1000 Res 8.

Brunck, M.E., Andersen, S.B., Timmins, N.E., Osborne, G.W., and Nielsen, L.K. (2014). Absolute counting of neutrophils in whole blood using flow cytometry. Cytometry A 85, 1057–1064.

Cahalane, D.J., Charvet, C.J., and Finlay, B.L. (2014). Modeling local and cross-species neuron number variations in the cerebral cortex as arising from a common mechanism. Proc Natl Acad Sci U S A 111, 17642–17647.

Calegari, F., Haubensak, W., Haffner, C., and Huttner, W.B. (2005). Selective lengthening of the cell cycle in the neurogenic subpopulation of neural progenitor cells during mouse brain development. J Neurosci 25, 6533–6538.

Caviness, V.S., Jr., Takahashi, T., and Nowakowski, R.S. (1995). Numbers, time and neocortical neuronogenesis: a general developmental and evolutionary model. Trends Neurosci 18, 379–383.

Chao, H.W., Doi, M., Fustin, J.M., Chen, H., Murase, K., Maeda, Y., Hayashi, H., Tanaka, R., Sugawa, M., Mizukuchi, N., et al. (2017). Circadian clock regulates hepatic polyploidy by modulating Mkp1-Erk1/2 signaling pathway. Nat Commun 8, 2238.

Darzynkiewicz, Z., Halicka, H.D., and Zhao, H. (2010). Analysis of cellular DNA content by flow and laser scanning cytometry. Adv Exp Med Biol 676, 137–147.

Dominguez, M.H., Ayoub, A.E., and Rakic, P. (2013). POU-III transcription factors (Brn1, Brn2, and Oct6) influence neurogenesis, molecular identity, and migratory destination of upper-layer cells of the cerebral cortex. Cereb Cortex 23, 2632–2643.

Fei, J.F., Haffner, C., and Huttner, W.B. (2014). 3’ UTR-dependent, miR-92-mediated restriction of Tis21 expression maintains asymmetric neural stem cell division to ensure proper neocortex size. Cell Rep 7, 398–411.

Freret-Hodara, B., Cui, Y., Griveau, A., Vigier, L., Arai, Y., Touboul, J., and Pierani, A. (2017). Enhanced Abventricular Proliferation Compensates Cell Death in the Embryonic Cerebral Cortex. Cereb Cortex 27, 4701–4718.

Fulwyler, M.J. (1965). Electronic separation of biological cells by volume. Science 150, 910–911.

Gao, P., Postiglione, M.P., Krieger, T.G., Hernandez, L., Wang, C., Han, Z., Streicher, C., Papusheva, E., Insolera, R., Chugh, K., et al. (2014). Deterministic progenitor behavior and unitary production of neurons in the neocortex. Cell 159, 775–788.

Gasnereau, I., Ganier, O., Bourgain, F., de Gramont, A., Gendron, M.C., and Sobczak-Thepot, J. (2007). Flow cytometry to sort mammalian cells in cytokinesis. Cytometry A 71, 1–7.

Greig, L.C., Woodworth, M.B., Galazo, M.J., Padmanabhan, H., and Macklis, J.D. (2013). Molecular logic of neocortical projection neuron specification, development and diversity. Nat Rev Neurosci 14, 755–769.

Gusel’nikova, V.V., and Korzhevskiy, D.E. (2015). NeuN As a Neuronal Nuclear Antigen and Neuron Differentiation Marker. Acta Naturae 7, 42–47.

Haubensak, W., Attardo, A., Denk, W., and Huttner, W.B. (2004). Neurons arise in the basal neuroepithelium of the early mammalian telencephalon: a major site of neurogenesis. Proc Natl Acad Sci U S A 101, 3196–3201.

Herculano-Houzel, S., Watson, C., and Paxinos, G. (2013). Distribution of neurons in functional areas of the mouse cerebral cortex reveals quantitatively different cortical zones. Front Neuroanat 7, 35.

Jungas, T., Perchey, R.T., Fawal, M., Callot, C., Froment, C., Burlet-Schiltz, O., Besson, A., and Davy, A. (2016). Eph-mediated tyrosine phosphorylation of Citron Kinase controls abscission. J Cell Biol 214, 555–569.

Juric-Sekhar, G., and Hevner, R.F. (2019). Malformations of Cerebral Cortex Development: Molecules and Mechanisms. Annu Rev Pathol 14, 293–318.

Kischel, A., Audouard, C., Fawal, M.A., and Davy, A. (2020). Ephrin-B2 paces neuronal production in the developing neocortex. BMC Dev Biol 20, 12.

Knock, E., Pereira, J., Lombard, P.D., Dimond, A., Leaford, D., Livesey, F.J., and Hendrich, B. (2015). The methyl binding domain 3/nucleosome remodelling and deacetylase complex regulates neural cell fate determination and terminal differentiation in the cerebral cortex. Neural Dev 10, 13.

Kushchayev, S.V., Moskalenko, V.F., Wiener, P.C., Tsymbaliuk, V.I., Cherkasov, V.G., Dzyavulska, I.V., Kovalchuk, O.I., Sonntag, V.K., Spetzler, R.F., and Preul, M.C. (2012). The discovery of the pyramidal neurons: Vladimir Betz and a new era of neuroscience. Brain 135, 285–300.

Lanctot, A.A., Guo, Y., Le, Y., Edens, B.M., Nowakowski, R.S., and Feng, Y. (2017). Loss of Brap Results in Premature G1/S Phase Transition and Impeded Neural Progenitor Differentiation. Cell Rep 20, 1148–1160.

Llorca, A., Ciceri, G., Beattie, R., Wong, F.K., Diana, G., Serafeimidou-Pouliou, E., Fernandez-Otero, M., Streicher, C., Arnold, S.J., Meyer, M., et al. (2019). A stochastic framework of neurogenesis underlies the assembly of neocortical cytoarchitecture. Elife 8.

Lodato, S., Shetty, A.S., and Arlotta, P. (2015). Cerebral cortex assembly: generating and reprogramming projection neuron diversity. Trends Neurosci 38, 117–125.

Lopez-Sanchez, N., and Frade, J.M. (2013). Genetic evidence for p75NTR-dependent tetraploidy in cortical projection neurons from adult mice. J Neurosci 33, 7488–7500.

Lui, J.H., Hansen, D.V., and Kriegstein, A.R. (2011). Development and evolution of the human neocortex. Cell 146, 18–36.

Malatesta, P., Hartfuss, E., and Gotz, M. (2000). Isolation of radial glial cells by fluorescent-activated cell sorting reveals a neuronal lineage. Development 127, 5253–5263.

McKinnon, K.M. (2018). Flow Cytometry: An Overview. Curr Protoc Immunol 120, 5 1 1–5 1 11.

Mihalas, A.B., and Hevner, R.F. (2018). Clonal analysis reveals laminar fate multipotency and daughter cell apoptosis of mouse cortical intermediate progenitors. Development 145.

Miller, D.J., Bhaduri, A., Sestan, N., and Kriegstein, A. (2019). Shared and derived features of cellular diversity in the human cerebral cortex. Curr Opin Neurobiol 56, 117–124.

Miyata, T., Kawaguchi, A., Okano, H., and Ogawa, M. (2001). Asymmetric inheritance of radial glial fibers by cortical neurons. Neuron 31, 727–741.

Noctor, S.C., Flint, A.C., Weissman, T.A., Dammerman, R.S., and Kriegstein, A.R. (2001). Neurons derived from radial glial cells establish radial units in neocortex. Nature 409, 714–720.

Ou, F., McGoverin, C., Swift, S., and Vanholsbeeck, F. (2017). Absolute bacterial cell enumeration using flow cytometry. J Appl Microbiol 123, 464–477.

Paredes, M.F., Sorrells, S.F., Garcia-Verdugo, J.M., and Alvarez-Buylla, A. (2016). Brain size and limits to adult neurogenesis. J Comp Neurol 524, 646–664.

Picco, N., Garcia-Moreno, F., Maini, P.K., Woolley, T.E., and Molnar, Z. (2018). Mathematical Modeling of Cortical Neurogenesis Reveals that the Founder Population does not Necessarily Scale with Neurogenic Output. Cereb Cortex 28, 2540–2550.

Postel, M., Karam, A., Pezeron, G., Schneider-Maunoury, S., and Clement, F. (2019). A multiscale mathematical model of cell dynamics during neurogenesis in the mouse cerebral cortex. BMC Bioinformatics 20, 470.

Sahara, S., Kodama, T., and Stevens, C.F. (2020). A common rule governing differentiation kinetics of mouse cortical progenitors. Proc Natl Acad Sci U S A.

Sessa, A., Mao, C.A., Hadjantonakis, A.K., Klein, W.H., and Broccoli, V. (2008). Tbr2 directs conversion of radial glia into basal precursors and guides neuronal amplification by indirect neurogenesis in the developing neocortex. Neuron 60, 56–69.

Sigl-Glockner, J., and Brecht, M. (2017). Polyploidy and the Cellular and Areal Diversity of Rat Cortical Layer 5 Pyramidal Neurons. Cell Rep 20, 2575–2583.

Telley, L., Agirman, G., Prados, J., Amberg, N., Fievre, S., Oberst, P., Bartolini, G., Vitali, I., Cadilhac, C., Hippenmeyer, S., et al. (2019). Temporal patterning of apical progenitors and their daughter neurons in the developing neocortex. Science 364.

Telley, L., Govindan, S., Prados, J., Stevant, I., Nef, S., Dermitzakis, E., Dayer, A., and Jabaudon, D. (2016). Sequential transcriptional waves direct the differentiation of newborn neurons in the mouse neocortex. Science 351, 1443–1446.

Wang, L., Hou, S., and Han, Y.G. (2016). Hedgehog signaling promotes basal progenitor expansion and the growth and folding of the neocortex. Nat Neurosci 19, 888–896.

Westra, J.W., Barral, S., and Chun, J. (2009). A reevaluation of tetraploidy in the Alzheimer’s disease brain. Neurodegener Dis 6, 221–229.

Woodworth, M.B., Greig, L.C., Kriegstein, A.R., and Macklis, J.D. (2012). SnapShot: cortical development. Cell 151, 918–918 e911.

Yang, Y., Geldmacher, D.S., and Herrup, K. (2001). DNA replication precedes neuronal cell death in Alzheimer’s disease. J Neurosci 21, 2661–2668.

Yoon, K.J., Ringeling, F.R., Vissers, C., Jacob, F., Pokrass, M., Jimenez-Cyrus, D., Su, Y., Kim, N.S., Zhu, Y., Zheng, L., et al. (2017). Temporal Control of Mammalian Cortical Neurogenesis by m(6)A Methylation. Cell 171, 877–889 e817.

Young, N.A., Flaherty, D.K., Airey, D.C., Varlan, P., Aworunse, F., Kaas, J.H., and Collins, C.E. (2012). Use of flow cytometry for high-throughput cell population estimates in brain tissue. Front Neuroanat 6, 27.

