## Supplemental Figures and Table for "Population Dynamics and Neuronal Polyploidy in the Developing Neocortex"

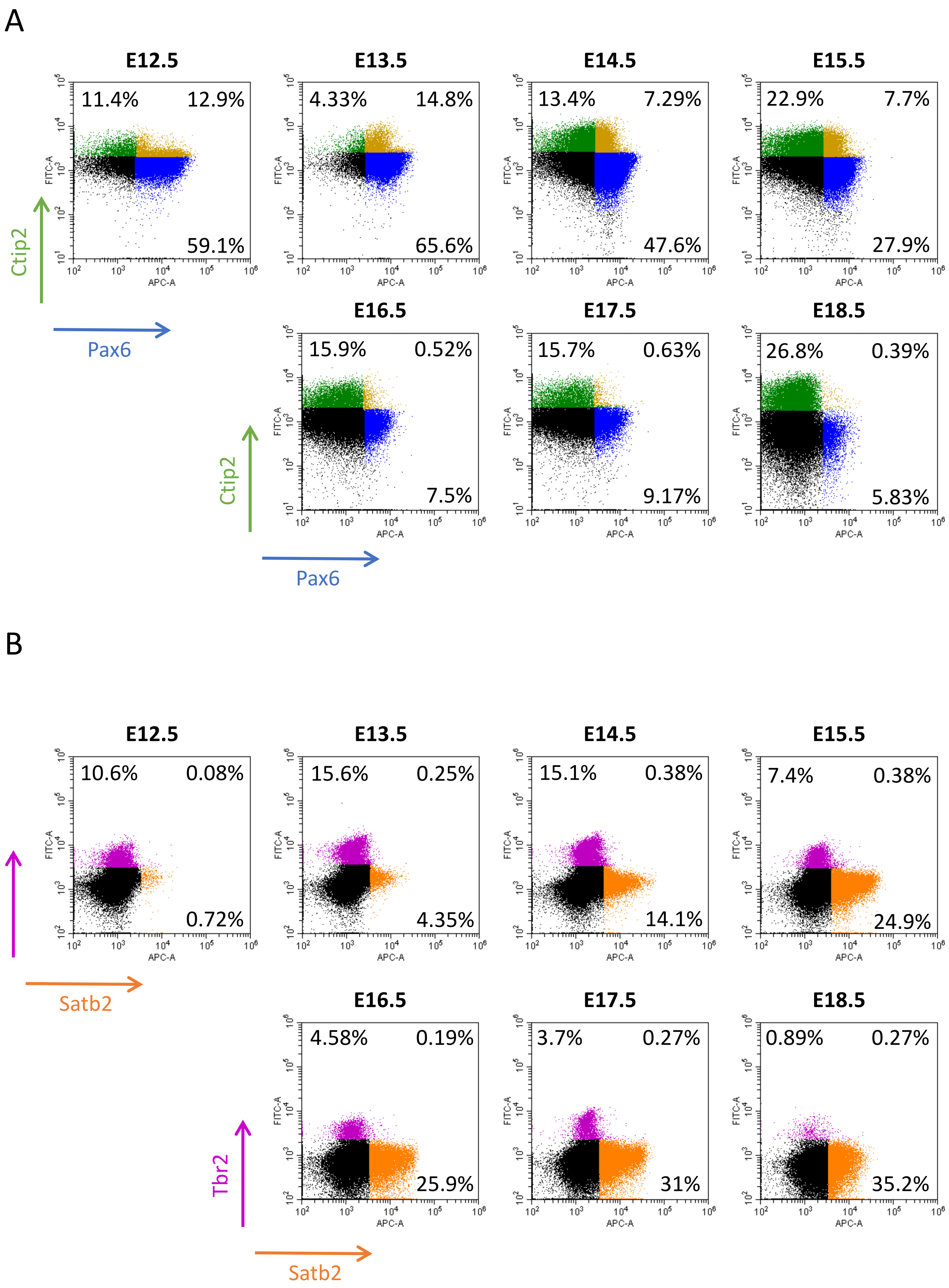


**Sup Figure 1: Detailed analysis of Pax6/Ctip2 and Tbr2/Satb2 populations during neocortex development**

**Sup Figure 1: Detailed analysis of Pax6/Ctip2 and Tbr2/Satb2 double positive populations during neocortex development (Related to Figure 2).**

A: Representative dot plots for each developmental stage of Ctip2-Alexa fluor488 associated fluorescence intensity (vertical axis) versus Pax6-Alexa fluor647 associated fluorescence intensity (horizontal axis). Blue dots represent Pax6+ cells. Green dots represent Ctip2+ cells. Yellow dots represent double positive for Ctip2 and Pax6 (blue, upper right quadrant). The fraction of single and double positive cells is indicated in each quadrant.

B: Representative dot plots for each developmental stage of Tbr2-Alexa fluor488 associated fluorescence intensity (vertical axis) versus Satb2-Alexa fluor647 associated fluorescence intensity (horizontal axis). Purple dots represent Tbr2+ cells. Orange dots represent Satb2+ cells. The fraction of single and double positive cells is indicated in each quadrant.

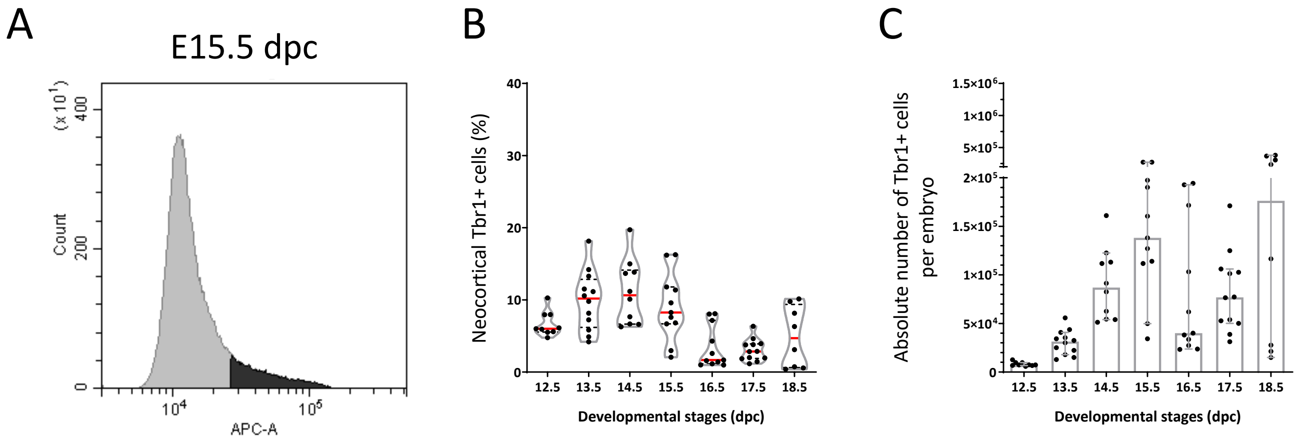


**Sup Figure 2: Analysis of Tbr1+ cell population during neocortex development**

**Sup Figure 2: Analysis of Tbr1+ cell population during neocortex development (Related to Figure 2 and Figure 3).**

A: Histogram plot of cells from a E15.5 embryo showing Tbr1 associated Alexafluor 647 signal intensity detected on APC-A channel (Tbr1+ cells are in black).

B: Relative proportion of Tbr1+ cells at each indicated developmental stage. Each dot represents one embryo. Violin representation of the data displays the distribution (shape); the median (red line) and $1^{st}$ and $3^{rd}$ quartile (dotted black lines).

C: Absolute number of Tbr1+ cells in the neocortex per embryo at each developmental stage. The histogram bars correspond to the mean with 95% confidence interval error bars. Each dot represents the calculated value for one embryo.

A

| sample id | age | PI volume | beads volume | lot beads concentration | acquired beads | acquired cells | Total cell number |
|---|---|---|---|---|---|---|---|
| P300-1 | 300 days | 400 µl | 50 µl | 1 019 beads / µl | 1 317 | 63 687 | 24 589 855 |
| P300-2 | 300 days | 400 µl | 50 µl | 1 019 beads / µl | 1 498 | 63 019 | 21 391 964 |

B

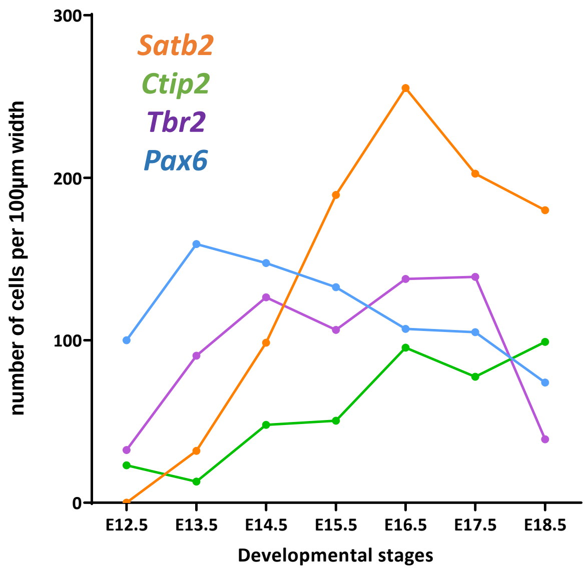


**Sup figure 3: Comparison with published data at post-natal and pre-natal stages**

**Sup Figure 3: Comparison with published data at post-natal and pre-natal stages (Related to Figure 3).**

A: Raw data and calculation with the bead-based methods of the absolute number of cells for 2 distinct cortical samples from adult mice at 300 days post natal.

B: Comparative dynamics of the 4 cell populations over the course of neocortex development: Pax6+ (blue), Tbr2+ (purple), Ctip2+ (green), Satb2+ (orange). 2D data was collected from a bibliographic survey (Supplementary Table 2). Dots represent mean values for each developmental stage.

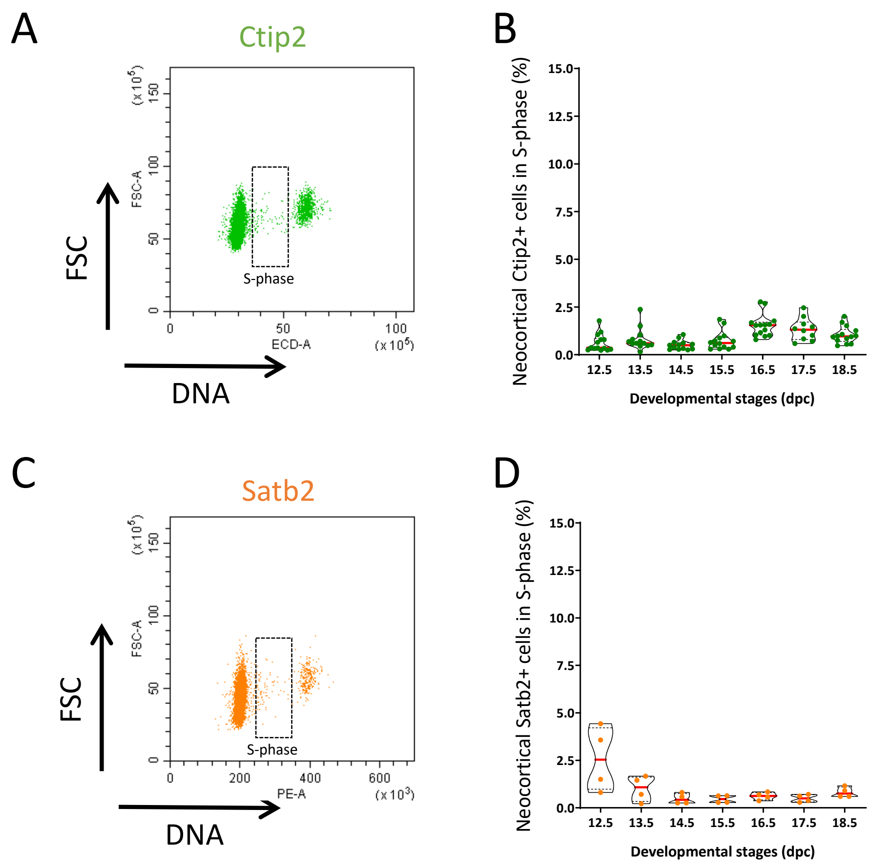


**Sup Figure 4: Cell cycle parameters for the 2 subtypes of cortical neurons over the course of neocortex development**

**Sup Figure 4: Cell cycle parameters for the 2 subtypes of cortical neurons over the course of neocortex development (Related to Figure 4).**

A: Example of a dot plot of DNA content for Ctip2+ cells estimated by Propidium Iodide incorporation (horizontal axis) versus FSC relative cell size (vertical axis) for one embryo. Cells in S-phase are highlighted.

B: Distribution of the proportion of Ctip2+ cells in S-phase at each developmental stage. Each dot represents one embryo. Violin representation of the data displays the distribution (shape); the median (red line) and $1^{st}$ and $3^{rd}$ quartile (dotted black lines).

C: Example of a dot plot of DNA content for Satb2+ cells estimated by Propidium Iodide incorporation (horizontal axis) versus FSC relative cell size (vertical axis) for one embryo. Cells in S-phase are highlighted.

B: Distribution of the proportion of Satb2+ cells in S-phase at each developmental stage. Each dot represents one embryo. Violin representation of the data displays the distribution (shape); the median (red line) and $1^{st}$ and $3^{rd}$ quartile (dotted black lines).

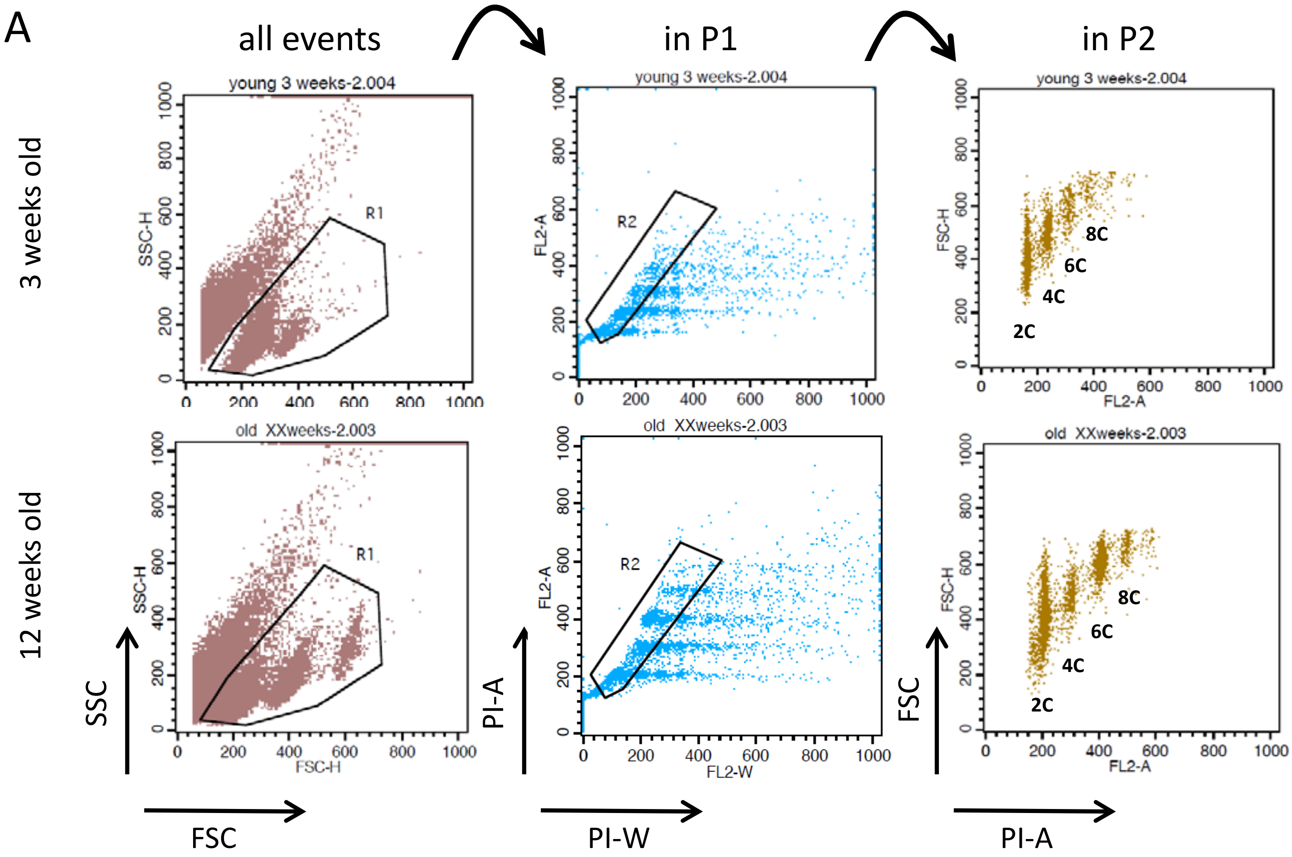


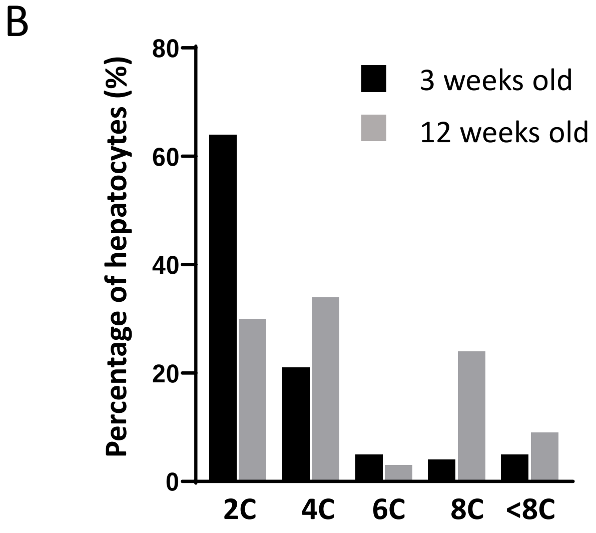


**Sup Figure 5: Quantification of polyploid cells in adult liver using the flow cytometry procedure**

**Sup Figure 5: Quantification of polyploid cells in adult liver using the flow cytometry procedure (Related to Figure 5).**

A: Side scatter (SSC) versus Forward Scatter (FSC) dot plot of all acquired events showing the gate R1 (left). Dot plot for Propidium Iodide Width signal intensity (PI-W) versus Propidium Iodide Area signal intensity (PI-A) of R1 gated events showing the gate R2 corresponding to the singlets retained for further analysis (middle). R1 and R2 gated cells plotted with FSC on vertical axis and PI fluorescence signal intensity (PI-H) on horizontal axis (right).

B: Fraction of hepatocytes (%) according to their DNA content. Bars represent one measurement of one sample.

Supplemental Table 3. Related to Figure 3.

| stage | Pax6 | Tbr2 | Ctip2 | Satb2 | width (um) | Reference |
|---|---|---|---|---|---|---|
| E10.5 | 65 | 10 | | | 100 | Fei et al. Cell Reports |
| E11 | 120 | 16 | | | 100 | Alsio et al. PNAS |
| E12.5 | 80 | 26 | | | 100 | Lanctot et al. Cell Reports |
| E12.5 | 120 | 39 | 23 | | 100 | Postel et al. BMC Bioinfo |
| E13 | 190 | 56 | | | 100 | Alsio et al. PNAS |
| E13.5 | 110 | 120 | | 21 | 100 | Kischel et al. BMC Dev Biol |
| E13.5 | 182 | 80 | | | 100 | Fei et al. Cell Reports |
| E13.5 | 155 | 106 | 13 | 43 | 100 | Postel et al. BMC Bioinfo |
| E14.5 | 150 | 120 | | 65 | 100 | Kischel et al. BMC Dev Biol |
| E14.5 | 145 | 133 | 48 | 132 | 100 | Postel et al. BMC Bioinfo |
| E15 | 160 | 68 | | | 100 | Alsio et al. PNAS |
| E15 | 124 | 109 | 38 | 146 | 100 | Postel et al. BMC Bioinfo |
| E15.5 | 114 | 142 | 63 | 233 | 100 | Postel et al. BMC Bioinfo |
| E16 | 104 | 161 | 86 | 230 | 100 | Postel et al. BMC Bioinfo |
| E16.5 | | | | 150 | 100 | Kischel et al. BMC Dev Biol |
| E16.5 | 69 | 130 | 105 | 251 | 100 | Postel et al. BMC Bioinfo |
| E16.5 | 80 | 150 | | | 100 | Wang et al. Nature Neurosci. |
| E16.5 | 175 | 110 | | | 100 | Fei et al. Cell Reports |
| E17 | 90 | | | | 100 | Alsio et al. PNAS |
| E17.5 | | | 116 | 162 | 100 | Alsio et al. PNAS |
| E17.5 | 95 | 128 | 39 | 243 | 100 | Postel et al. BMC Bioinfo |
| E17.5 | 130 | 150 | | | 100 | Yoon et al. Cell |
| E18.5 | | | | 180 | 100 | Kischel et al. BMC Dev Biol |
| E18.5 | 74 | 39 | 99 | 180 | 100 | Postel et al. BMC Bioinfo |
